# Biopsychosocial Correlates of Resting and Stress-Reactive Salivary GDF15: Preliminary Findings

**DOI:** 10.1101/2025.02.27.640377

**Authors:** Cynthia C. Liu, Caroline Trumpff, Qiuhan Huang, Robert-Paul Juster, Martin Picard

## Abstract

Growth differentiation factor 15 (GDF15) is a biomarker of energetic stress related to aging, disease, and mitochondrial defects. We recently showed that GDF15 is quantifiable in saliva and acutely inducible by psychosocial stress. To date, the associations between GDF15 and biopsychosocial factors and individual characteristics remain unknown. Here, in a sample of healthy working adults (*n* = 198, 70% females), we first confirmed that salivary GDF15 reacts to acute psychosocial stress, peaking 10 min following a socio-evaluative stress paradigm (+28.3%, *g* = 0.50, *p* < 0.0001). We then explored associations between i) baseline GDF15 and ii) GDF15 stress reactivity and a variety of trait- and state-level biopsychosocial factors including sex and gender characteristics; measures of mental health, stress, and burnout; physical health and health behaviors; and anthropometric and blood-based metabolic biomarkers. Baseline salivary GDF15 was higher in men than in women and was positively correlated with testosterone, while negatively correlated with estrogen and traditionally feminine gender roles. Of the psychosocial factors examined, we found that work-related stress variables were most consistently related to GDF15, with work-related cynicism, burnout, and emotional exhaustion predicting higher GDF15 reactivity, while job-related autonomy and utilization of competence predicted smaller GDF15 responses. Consistent with GDF15’s induction in metabolic and renal diseases, baseline GDF15 was also positively correlated with indirect markers of metabolic disease including waist-to-hip ratio, creatinine, and albumin. Finally, participants with greater GDF15 reactivity also exhibited greater cortisol reactivity, consistent with the role of GDF15 in stress regulation and energy mobilization. Together, this exploratory analysis of salivary GDF15 suggest new biological and psychosocial correlates, calling for large-scale studies connecting human experiences with biological markers of energetic stress.

## 1. Introduction

The cytokine/metabokine *growth differentiation factor 15* (GDF15) is the most prominently elevated circulating protein across several medical conditions, including cancer, cardiovascular, kidney, Alzheimer’s, autoimmune, and rare mitochondrial diseases (Welsh, Sapinoso et al. 2003, Amstad, Coray et al. 2020, Lin, Ji et al. 2020, Sharma, Reinstadler et al. 2021, He, de Souto Barreto et al. 2022, Li, Li et al. 2022, Gadd, Hillary et al. 2024, Guo, You et al. 2024, Lyu, Xv et al. 2024, Tang, Liu et al. 2024). GDF15 is a leading biomarker of aging—increasing by 100-150% on average per decade of life (Tanaka, Biancotto et al. 2018, Lehallier, Gate et al. 2019, Johnson, Shokhirev et al. 2020, Cefis, Marcangeli et al. 2025, Trumpff, Huang et al. 2025). Blood GDF15 is also dramatically elevated during pregnancy, with increases up to 200-fold (Andersson-Hall, Joelsson et al. 2020), which in extreme cases causes excessive nausea and vomiting (i.e., hyperemesis gravidarum (Fejzo, Rocha et al. 2024)). Beyond these cross-sectional associations establishing GDF15 as a critical marker of disease, GDF15 also predicts the incidence of neurological, cardiac, metabolic, and psychiatric disorders over 1-2 decades, as well as all-cause mortality (Gadd, Hillary et al. 2024, Deng, You et al. 2025). Together, these findings highlight the clinical and research importance of understanding the psychobiological factors that regulate GDF15 biology.

We recently showed that, in addition to blood, GDF15 can be measured in human saliva, where it appears to dynamically respond to acute psychosocial stress (Huang, Monzel et al. 2024). A study of 70 healthy individuals showed that, much like cortisol and other stress mediators, salivary GDF15 increased by an average of ∼43% within 5 minutes of psychosocial stress induction (i.e., a speech delivery task).

Interestingly, the relative magnitude of plasma GDF15 response was blunted in people with mitochondrial disorders, though not in saliva. These individuals have OxPhos defects that trigger cellular energetic deficiency, contributing to systemically elevated baseline GDF15 levels (Huang, Trumpff et al. 2024). People with psychiatric disorders (Frye, Nassan et al. 2015, Kumar, Millischer et al. 2017, Yang, Barbosa et al. 2018, Mastrobattista, Lenze et al. 2023, Pan, Naviaux et al. 2023) and comorbid mental-physical disorders (Lu, Duan et al. 2020, Sun, Peng et al. 2021, Zang, Zhu et al. 2022, Li, Mei et al. 2023) also exhibit elevated blood GDF15. Together, these new data suggest two things. First, salivary GDF15 may be an accessible marker of stress-reactivity and metabolic or mitochondrial health status. Second, mental and metabolic signaling pathways may converge in the regulation of GDF15 signaling.

Beyond acute laboratory stressors and clinical diagnoses, less is known about how GDF15 relates to sociodemographic, psychosocial, and health-related behaviors. Initial studies have investigated associations between circulating GDF15 levels and education, income, and poverty levels—finding that GDF15 was negatively correlated with years of education and household income (Shafi, Bøttcher et al. 2022).

Interestingly, the risk of all-cause mortality associated with GDF15 was lower in those below the poverty cutoff (Freeman, Noren Hooten et al. 2020, Shafi, Bøttcher et al. 2022). In some studies, GDF15 was also related to lifestyle habits, exhibiting negative correlations with healthy dietary patterns, but positive correlations with alcohol consumption and smoking (Krintus, Braga et al. 2019, Ortolá, García-Esquinas et al. 2021, Ortolá, García-Esquinas et al. 2022). These initial results paint a picture that suggests adverse exposures may trigger energetically costly allostatic or stress responses (Bobba-Alves, Juster et al. 2022), while health-promoting behaviors may relieve the organism of the cost of stress (Bobba-Alves, Sturm et al. 2023), possibly dialing the GDF15 axis up and down, respectively. The role of GDF15 may then be to convey energetic stress to the brain to promote adaptation and survival (Lockhart, Saudek et al. 2020, Monzel, Levin et al. 2024). However, further research is needed to understand the relationship of salivary GDF15 with a broad range of state- and trait-level biopsychosocial factors, including the magnitude and directionality of relationships between GDF15 and potential risk and resilience factors.

Further research is also needed to clarify sex differences in GDF15 and the relationships between GDF15 and sex hormones in healthy populations. In the context of clinically diagnosed disorders, blood GDF15 was found to be inversely correlated with testosterone levels in males with sex-hormone imbalances, coronary heart disease, HBV-associated hepatocellular carcinoma, and major depressive disorder (Liu, Dai et al. 2019, Chen, Dai et al. 2020, Peng, Li et al. 2021, Sun, Peng et al. 2021, Li, Mei et al. 2023, Pan, Naviaux et al. 2023). In females, blood GDF15 was inversely correlated with estrogen during menopausal hormone therapy (Faubion, White et al. 2020). While these findings suggest some sex-specificity in GDF15 biology, no work has been done on *salivary* GDF15 and sex or sociocultural gender (Barr, Popkin et al. 2024) characteristics in healthy individuals.

Here, we begin to address some of these knowledge gaps by examining salivary GDF15 reactivity to the Trier Social Stress Test (TSST) in healthy adults assessed in a study of psychiatric hospital workers (Juster, Raymond et al. 2016). This study of stress physiology and mental health provided associations with sex hormones (Juster, Raymond et al. 2016), gender roles (Juster, Pruessner et al. 2016), and occupational gender roles (Kerr, Barbosa Da Torre et al. 2021), among other constructs. Using a discovery approach emphasizing patterns of associations over the statistical significance of specific tests, we explore relationships between both i) baseline GDF15 and ii) GDF15 reactivity with a variety of biopsychosocial factors, including sex and gender characteristics, mental and physical health variables, and anthropometric and blood-based metabolic biomarkers, together with the magnitude of stress reactivity among other psychobiological parameters. With these preliminary findings, we hope to lay the foundation for additional research exploring the role of GDF15 around psychobiological processes linking energetic stress and human experiences.

## 2. Materials and Methods

### 2.1 Participants

Participants were psychiatric hospital workers from a large psychiatric hospital in Canada, who were recruited as part of a larger study (Juster, Raymond et al. 2016). The study was approved by the ethics board of *Institut universitaire en santé mentale de Montréal* and adheres to the Declaration of Helsinki. A total of 204 participants were recruited (*n* = 144 female, *n* = 60 male), with female participants of varying reproductive statuses (cycling: *n* = 55; contraceptive: *n* = 47; post-menopausal: *n* = 42). All participants read and signed the consent form. Specific recruitment criteria and sample characteristics have been previously reported (Juster, Raymond et al. 2016). Briefly, participants were screened for history of known cardiovascular disease, neurological disorder, head trauma, thyroid problems, and diabetes mellitus, and exclusion criteria included use of antiseizure, antihypertensive, anxiolytics, or antidepressant medications.

### 2.2 Procedure

Participants visited the laboratory twice. The first visit involved informed consent, completion of a cognitive task (Dumont, Marin et al. 2020), a Trier Social Stress Test (TSST) (Kirschbaum, Pirke et al. 2008), and a debriefing session. Anticipatory stress was assessed 10 minutes before the TSST. To assess cortisol and GDF15 reactivity, 2mL of saliva was collected via sterilized 5 ml 57 × 15.3 mm screw cap tubes (Sarstedt^®^, Item No. 62.558.201) guided with thick straws at six timepoints before, during, and after the TSST (−10, 0, 10, 20, 30, 40min). Blood pressure and heart rate were also measured at these time points. An additional saliva sample was collected 20 minutes before the TSST to assess sex hormone levels. Between visits, participants completed questionnaires electronically. During the second visit, participants returned saliva samples they had collected at home and completed a physical examination and a blood draw. Physiological parameters and blood markers were used to assess allostatic load biomarkers as described previously (McEwen and Stellar 1993, Seeman, McEwen et al. 2001, Juster, McEwen et al. 2010, Picard, Juster et al. 2014).

### 2.3 Self-report measures

The 16-item Primary Appraisal Secondary Appraisal (PASA) scale (Gaab 2009) was administered 10 minutes prior to administration of the TSST to measure how participants perceived the stressful situation. This scale contains the primary appraisal subscales threat and challenge, the secondary appraisal subscales self-concept of competence and control expectancy, and the tertiary appraisal total score representing global anticipatory distress. Items on each subscale were rated from 1-6, with total scores ranging from 4-24. Higher scores indicate greater threat/challenge, competence/control, or distress.

Participants then completed a questionnaire package at home that included demographic questions, (i.e., age, sex, and years of education), questions about health-related behaviors (i.e., smoking, alcohol and caffeine consumption habits) and health status (i.e., number of mental and physical health condition), as well as a series of standardized psychosocial questionnaires. The subset of scales used in this analysis are described below, and the full set of questionnaires are described elsewhere (Juster and Lupien 2012, Juster, Raymond et al. 2016).

Gender roles were measured via the 60-item Bem Sex Role Inventory (BSRI) (Bem 1974, Bem 1978), which uses a 7-point Likert scale to measure traditionally defined masculine and feminine items. Masculine and feminine subscales were dichotomized to create four gender role classifications—feminine, masculine, androgynous, and undifferentiated—using the medians for each subscale (masculinity subscale: 4.6 for females and 4.8 for males, femininity subscale: 6.1 for females and 5.9 for males). Individuals that scored above the median on the femininity scale but below the median on the masculinity scale were deemed feminine. Individuals that scored above the median on the masculinity scale but below the median on the femininity scale were deemed masculine. Individuals that scored above the median on both scales were deemed androgynous. Individuals that scored below the median on both scales were deemed undifferentiated (Juster and Lupien 2012).

The Big Five personality traits were assessed via the Ten-item Personality Index (TIPI) (Gosling, Rentfrow et al. 2003). Participants responded to items on a scale of 1-7, with two items corresponding to each of the five personality traits (extraversion, agreeableness, conscientiousness, emotional stability, openness to experiences). Subscale totals range from 2-14, with higher scores indicating higher levels of the given personality trait.

Perceived social support was assessed via the 20-item Perceived Social Support Scale (Zimet, Dahlem et al. 1988). Participants responded to items on a scale of 1-5. Total scores range from 20-100, with higher scores representing greater levels of social support.

Self-Esteem was assessed using the 10-item Rosenberg Self-Esteem Scale (Rosenberg 1965). Participants responded to items on a scale of 0-3. Total scores range from 0-30, with higher scores representing higher self-esteem.

Emotional, psychological, and social well-being was assessed using the 14-item Mental Health Continuum scale (Keyes 2009). Participants responded to items on a scale of 0-5. Total scores range from 0-70, with higher scores representing greater levels of wellbeing.

Depressive symptoms were assessed using the 21-item Beck Depression Inventory II (Beck, Steer et al. 1996). Participants responded to items on a scale of 0-3. Total scores range from 0-63, with higher scores representing greater levels of depression.

Post-traumatic stress disorder (PTSD) symptoms were assessed using the PTSD Civilian Checklist (Blevins, Weathers et al. 2015). Participants responded to items on a scale of 1-5. Total scores range from 17-85, with higher scores representing more PTSD symptoms.

Memory complaints were assessed using the 6-item Memory Assessment Clinic-questionnaire (Crook, Feher et al. 1992). Participants responded to items on a scale of 1-5. Scores range from 7-35, with higher scores reflecting greater memory complaints.

Chronic stress was assessed using the English version of the 30-item Trier Inventory for the assessment of Chronic Stress (TICS) (Schulz 1995). Participants responded to items on a scale of 0-4. Resultant scores include a total score ranging from 0-120, as well as sub-scales ranging from 0-12 that include: work overload, work discontent, overextended at work, performance pressure at work, worry propensity, social overload, social tension, lack of social recognition, performance pressure in social interactions, and social isolation. Higher scores indicate greater stress.

Workplace stressors were assessed using the 26-item Job Content Questionnaire (JCQ) (Karasek, Brisson et al. 1998), which measured the content of participants work, include decision latitude (9-items) comprising skill utilization and work constraints, psychological demands (9-items), and social support from colleagues and supervisors (8-items). Participants responded to items on a scale of 1-4, with higher scores indicating more decision latitude, psychological demands, or social support, respectively.

Workplace characteristics were further assessed using the 16-item Effort-reward imbalance inventory questionnaire (ER) (Siegrist, Li et al. 2014), comprised of 6 items measuring effort and 10 items measuring reward, with subscales of esteem, promotion and security. Overcommitment was also measured by an additional 6 items. Participants responded to items on a scale of 1-4. In our data, the effort scale ranged from 6-29, the reward scale ranged from 0-39, and the overcommitment scale ranged from 0-0.9, with higher scores indicating more effort, reward, or overcommitment, respectively.

Burnout was measured using the 16-item Maslach Burnout Inventory (MBI) (Maslach 1996, Schaufeli, Leiter et al. 1996), which assesses the frequency of burnout symptoms. Participants respond to items on a scale of 0-6. Average scores were calculated for the three subscales: cynicism (5 items, scores range from 0-6), emotional exhaustion (5 items, scores range from 0-5.2), and professional efficacy (6 items, scores range from 1-6). We also calculated a total score with all subscales (scores range from 6-51.3) and a ratio score calculated as (emotional exhaustion + cynicism)/(professional efficacy) (scores range from 0-6.25). Higher scores indicate greater levels of burnout for all scores except for professional efficacy, where greater scores indicate less burnout.

Occupational Status Scores were assessed using the Nam-Powers-Boyd method applied to the 2001 National Occupational Classification for Statistics system (Nam 2000, Nam and Boyd 2004, Boyd 2008). Scores were calculated using Appendix A from (Boyd 2008) based on the median education and income from the Canadian census. Occupational prestige was ranked for 520 occupations (1 = Specialist Physicians to 520 = Trappers & Hunters) and education and income for each occupation was estimated. Sleep quality was assessed using the Pittsburgh Sleep Quality Index (PSQI) (Buysse, Reynolds et al. 1989), where participants scored each item from 0-3. Total scores range from 0-21, with higher scores indicating greater sleep difficulties.

### 2.4 Biological Measures

#### 2.4.1 GDF15 Assays

Salivary GDF15 levels were quantified using a high-sensitivity ELISA kit (R&D Systems, DGD150, SGD150) per manufacturer instructions. Samples were loaded directly and not diluted for these assays. Final plates were read by absorbance at 450nm on a Spectramax M2 (Spectramax Pro 6, Molecular Devices), and concentrations were interpolated using a four-parameter logistic (4PL) model in GraphPad Prism (version 10.3.1) using a standard curve run on each plate. Three plasma reference samples were run on each plate, and their inter-assay coefficients of variation (CV) were monitored. All the samples from the time course for a particular individual were kept on the same plate. Each sample was run in duplicate, with duplicates on separate plates where possible.

Final concentrations were calculated as the average of these duplicates. If the CV between duplicates was larger than 15%, the sample was rerun. If the CVs for the rerun samples were lower than the original, they were used, if not, the original values were retained. When it was not possible to rerun (e.g., no sample left), both duplicates were included and averaged. Values below the minimum detection range (2.0pg/ml) were considered undetectable, reported as NA, and excluded from analyses. Six participants did not have any saliva samples collected for GDF15 analysis, leaving a total of 198 participants with GDF15 data. A total of 1178 samples (unique timepoint/participant combinations) were analyzed.

#### 2.4.2 Cortisol and Sex Hormones

Salivary cortisol and sex hormones were assayed at the Saliva Laboratory of the Centre for Studies on Human Stress (www.humanstress.ca) (Juster, Raymond et al. 2016). Frozen samples were thawed and centrifuged at 1500 × *g* (3000 rpm) for 15 min. High-sensitivity enzymeimmuneassays were used for cortisol (Salimetrics^®^, No. 1-3102), estradiol (Salimetrics^®^, No. 1-3702), and progesterone (Salimetrics^®^ No. 1-1502). Testosterone was determined by expanded-range enzymeimmuneassay (Salimetrics^®^, No. 1-2402).

#### 2.4.3 Bloodwork assays

To assay blood biomarkers, 44ml of fasting blood was extracted by a licensed nurse during the second laboratory visit. Serum dehydroepiandrosterone-sulphate, C-reactive protein, albumin, creatinine, glycosylated hemoglobin, total cholesterol, high-density lipoprotein, and triglycerides was assessed using certified clinical facilities at the Louis H Lafontaine Hospital. Serum insulin and fibrinogen levels were assayed at Maisonneuve-Rosemont Hospital (Montreal, Quebec, Canada). Plasma tumor necrosis factor α and interleukin-6 was assayed at the Laboratory of the Centre for Studies on Human Stress.

#### 2.4.4 Allostatic load

Allostatic load was calculated as described previously (Juster, Pruessner et al. 2016) using the following variables: diurnal cortisol, TNF-alpha, IL6, fibrinogen, c-reactive protein, glycosylated hemoglobin, insulin, dehydroepiandrosterone-sulphate, creatinine, albumin, triglycerides, high-density lipoprotein cholesterol, total cholesterol, waist-to-hip ratio, body mass index, heart rate, TSST diastolic and systolic area under the curve to ground (AUCg), and TSST cortisol AUCg. Sex-specific allostatic load indices were determined with cut-offs for males and females separately; clinical allostatic load was determined based on clinical reference ranges, and hybrid allostatic load was determined with a combination of group-based cut-offs (based on the entire sample’s cut-offs without regard for sex). More details regarding allostatic load calculations, including cut-offs, can be found in a previous report (Juster, Raymond et al. 2016)

### 2.5 Statistical analyses

Data preprocessing, correlations, and regressions were conducted in R (version 4.3.2). Age correction of GDF15 data was performed using a linear regression model, whereby residuals to the linear regression line of each participant were subtracted from the median GDF15 value of the entire sample. GDF15 reactivity was calculated as the percentage change in GDF15 levels at 10min relative to −10min.

Mixed effect modeling was used to assess changes in GDF15 across the TSST time course (One-way ANOVA). Tukey’s multiple comparison tests were used to compare changes in GDF15 at each time point to baseline. These analyses were conducted in GraphPad Prism (version 10.3.1).

Partial spearman correlations (*r_s_*) with age as a covariate were calculated to assess associations between non-age corrected GDF15 variables (baseline GDF15 and GDF15 reactivity) and biopsychosocial variables. FDR corrections were calculated using the Benjamini-Hochberg method for each cluster of analyses (corresponding to each figure) for baseline GDF15 and GDF15 reactivity separately. Unadjusted p-values are reported in the figure heatmaps, and adjusted p-values (*p_adj_*) can be found in **Supplemental Table 1**. Group differences in age-adjusted GDF15 variables were assessed using Mann-Whitney U tests for comparisons between two groups, and Kruskal-Wallis H Tests and Dunn’s multiple comparison tests were used for comparisons between multiple groups.

K-means clustering and principal component dimensionality reduction analyses were performed in R. K-means clustering was performed using all self-report questionnaire variables, and 3 clusters were formed. GDF15 variables were compared between the 3 clusters. All data points, with clusters visualized, were then projected onto a 2-dimensional PCA space, and PCA loadings were determined. All code can be found at https://github.com/mitopsychobio/2025_GDF15_SGA_Liu.

In the spirit of the discovery approach taken in the following analyses, we provide interpretation and contrasts with the literature in sequence of our results.

## 3. Results

### 3.1 Salivary GDF15 is responsive to psychosocial stress and correlated with age

Across the 198 participants from which we sampled saliva at 6 timepoints before and after the TSST, salivary GDF15 exhibited significant variation over the 50min time course (Fixed effect of Time: *F* _3.7, 734.3_ *=* 8.35, *p* < 0.0001; **Figure 1A, B**). Specifically, salivary GDF15 increased by 28.3% on average from baseline to 10min post-TSST (*p* < 0.0001, Tukey’s test; **Figure 1B**). This peak was followed by a decline to near-baseline levels 10 minutes later, confirming the short half-life of salivary GDF15 reactivity (Huang, Monzel et al. 2024). These findings confirm that psychosocial stress induced an increase in salivary GDF15 concentrations similar to that observed for cortisol in this sample (Juster, Raymond et al. 2016).

**Figure 1.**
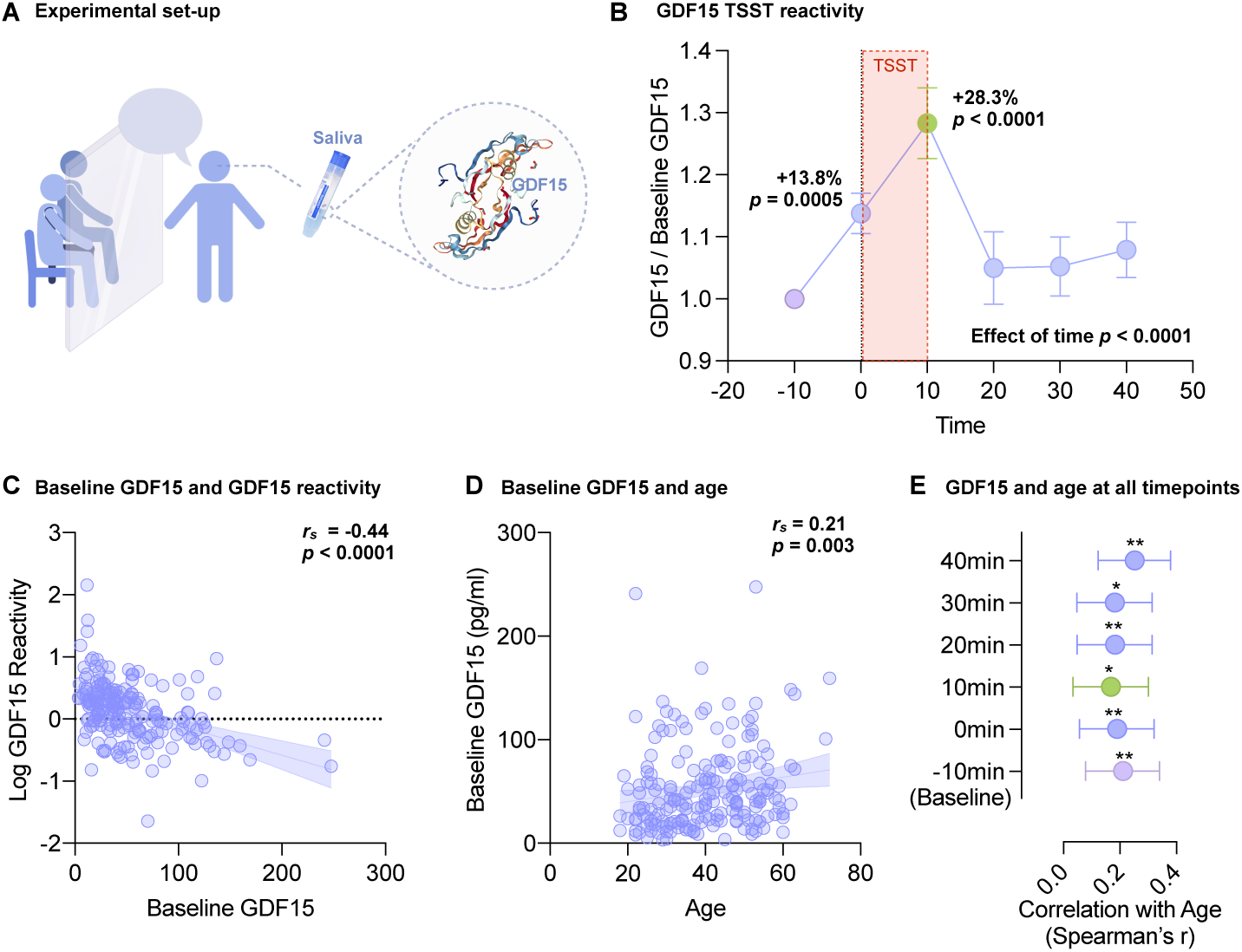
Salivary GDF15 is responsive to stress and correlated with age. **(A)** Experimental design measuring salivary GDF15 in response to psychosocial stress induction (TSST) (*n* = 194-197). **(B)** Average salivary GDF15 levels normalized to baseline (−10min) during the TSST task. Error bars represent SEM. Mixed-effect analysis revealed a significant effect of Time (*F*_(3.77, 734.3)_ = 8.35, *p* < 0.0001), with Tukey’s multiple comparisons test indicating significant differences between baseline, and 0min and 10min post-TSST. **(C)** Correlation between baseline GDF15 and log GDF15 reactivity (*n* = 196). **(D)** Correlation between baseline GDF15 and age (*n* = 197). **(E)** Forest plot of correlations between age and GDF15 at all time points (*n* = 194-198). Error bars represent 95% CI. Two-tailed significance indicated as \**p*<0.05, \*\**p*<0.01, \*\*\**p*<0.001, \*\*\*\**p*<0.0001. All correlations performed using Spearman’s rank correlation.

We then assessed whether this change in salivary GDF15 in response to psychosocial stress was correlated with baseline GDF15 concentrations. We defined GDF15 reactivity as the percent change in GDF15 from baseline to 10min post-TSST (where the average peak occurred). There was a moderate negative correlation between baseline GDF15 and GDF15 reactivity (*r_s_* = −0.44, *p* < 0.001; **Figure 1C**).

GDF15 measured in blood is a well-known marker of chronological age (Tanaka, Biancotto et al. 2018, Lehallier, Gate et al. 2019, Johnson, Shokhirev et al. 2020). In our sample, we found that salivary GDF15 was also elevated in older individuals, with small but significant positive correlations at baseline (*r_s_* = 0.21, *p* = 0.003; **Figure 1D**) and all other timepoints assessed (*r_s_*’s = 0.17 – 0.25, *p*’s = 0.0003 – 0.017; **Figure 1E**).

Replicating our prior study demonstrating the stress-inducibility of salivary GDF15 (Huang, Monzel et al. 2024) and the relationship between salivary GDF15 and age, we then moved to explore the relationships between GDF15—at baseline and in response to stress (reactivity)—and a variety of health-related biopsychosocial factors using a series of partial spearman correlations accounting for age.

### 3.2 Salivary GDF15 and sex and gender characteristics

Age-adjusted baseline salivary GDF15 was higher in males than in females (*z =* −4.17, *p* = 0.0002; **Figure 2A**). Moreover, salivary GDF15 differed between gender roles χ (3) = 10.9, *p* = 0.012), with multiple comparison tests indicating that individuals characterized as androgynous (i.e., possessing both high masculine and feminine personality traits) had lower levels of basal GDF15 than those characterized as feminine, masculine, or undifferentiated (i.e., low in both masculinity and femininity; **Table 1, Figure 2B**). As the Bem scale provides ranges for masculinity and femininity, we also assessed the correlations between salivary GDF15 and these continuous variables. Baseline GDF15 and femininity were negatively related (*r_s_*= −0.19, *p* = 0.010, *p_adj_*= 0.02), whereas masculinity trended in the opposite direction (*r_s_*= 0.13, *p* = 0.078, *p_adj_* = 0.12; **Figure 2C**). We also assessed correlations between our salivary GDF15 variables and concentrations of sex hormones (testosterone, estrogen, and progesterone) in both males and females. Baseline GDF15 was positively correlated with testosterone levels (*r_s_* = 0.25, *p* = 0.0004, *p_adj_*= 0.002) and negatively correlated with estrogen levels (*r_s_* = −0.19, *p* = 0.007, *p_adj_* = 0.02; **Figure 2D**). Together, these results (summarized in **Figure 2E**) suggest that salivary GDF15 levels may be lower in individuals who identified as female, have higher levels of estrogen, or who endorse behavioral characteristics traditionally deemed feminine, but higher in those who identify as male or who have higher levels of testosterone.

**Figure 2.**
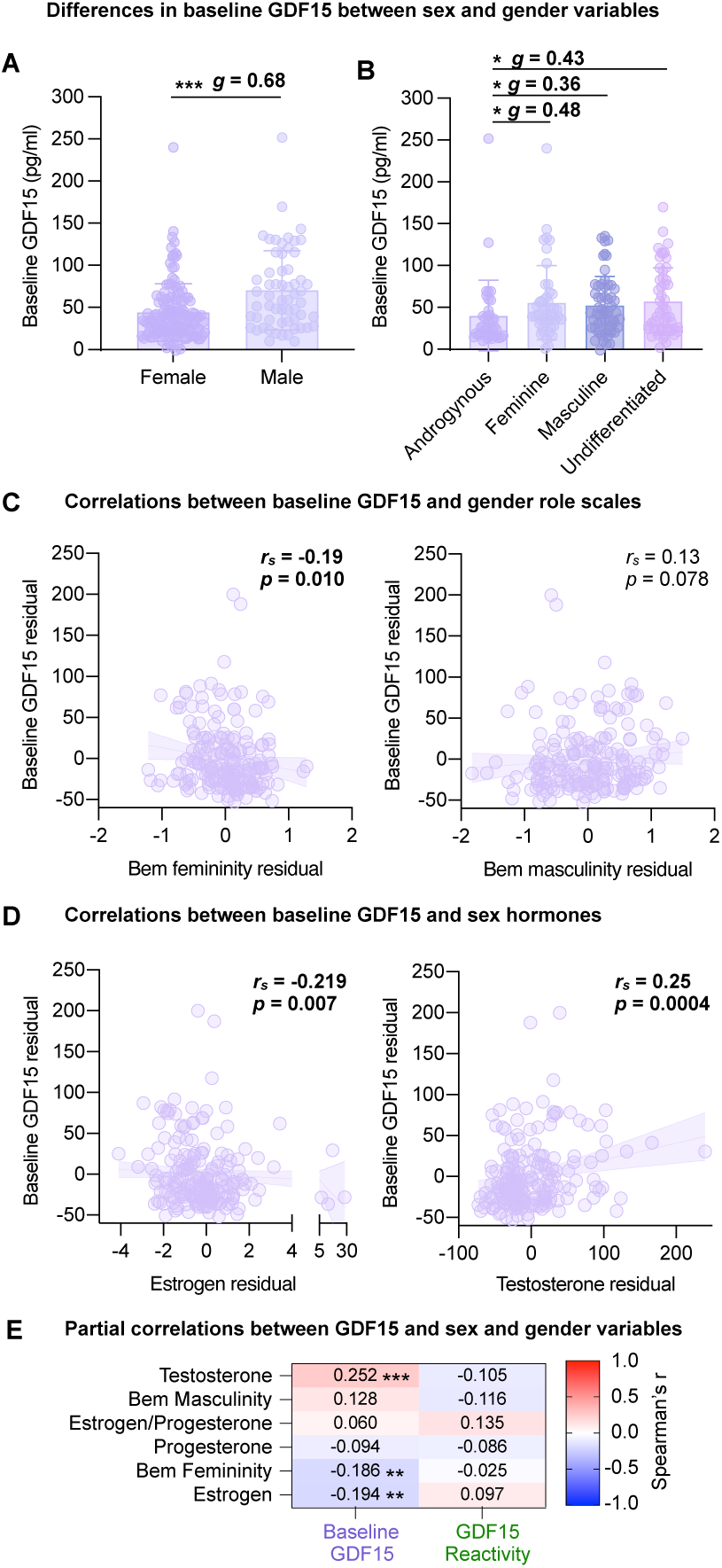
Salivary baseline GDF15 and GDF15 reactivity in correlations with sex and gender characteristics. **(A)** Difference in age-corrected baseline GDF15 between males (*n* = 59) and females (*n* = 138) tested with Mann-Whitney U test. Hedge’s *g* effect size labeled. **(B)** Differences in age-corrected baseline GDF15 between gender categories (Androgynous *n* = 37, Feminine *n* = 46, Masculine *n* = 58, Undifferentiated *n* = 52) as indicated by Dunn’s multiple comparison tests, with Hedge’s *g* effect size labeled. **(C)** Partial Spearman’s correlations between baseline GDF15 and masculinity (*n* = 193) or femininity (*n* = 193), adjusted for age. **(D)** Partial Spearman’s correlations between baseline GDF15 and testosterone (*n* = 197) or estrogen (*n* = 194), adjusted for age. **(E)** Heatmap showing partial correlations between sex and gender characteristics and baseline GDF15 or GDF15 reactivity (*n* = 53-197), adjusted for age. Sorted by descending effect size for baseline GDF15. Two-tailed significance indicated as \**p*<0.05, \*\**p*<0.01, \*\*\**p*<0.001, \*\*\*\**p*<0.0001. Unadjusted *p*-values are indicated in the figure; FDR-adjusted *p*-values can be found in Supplemental Table 1. All correlations performed using Spearman’s rank correlation.

**Table 1.**
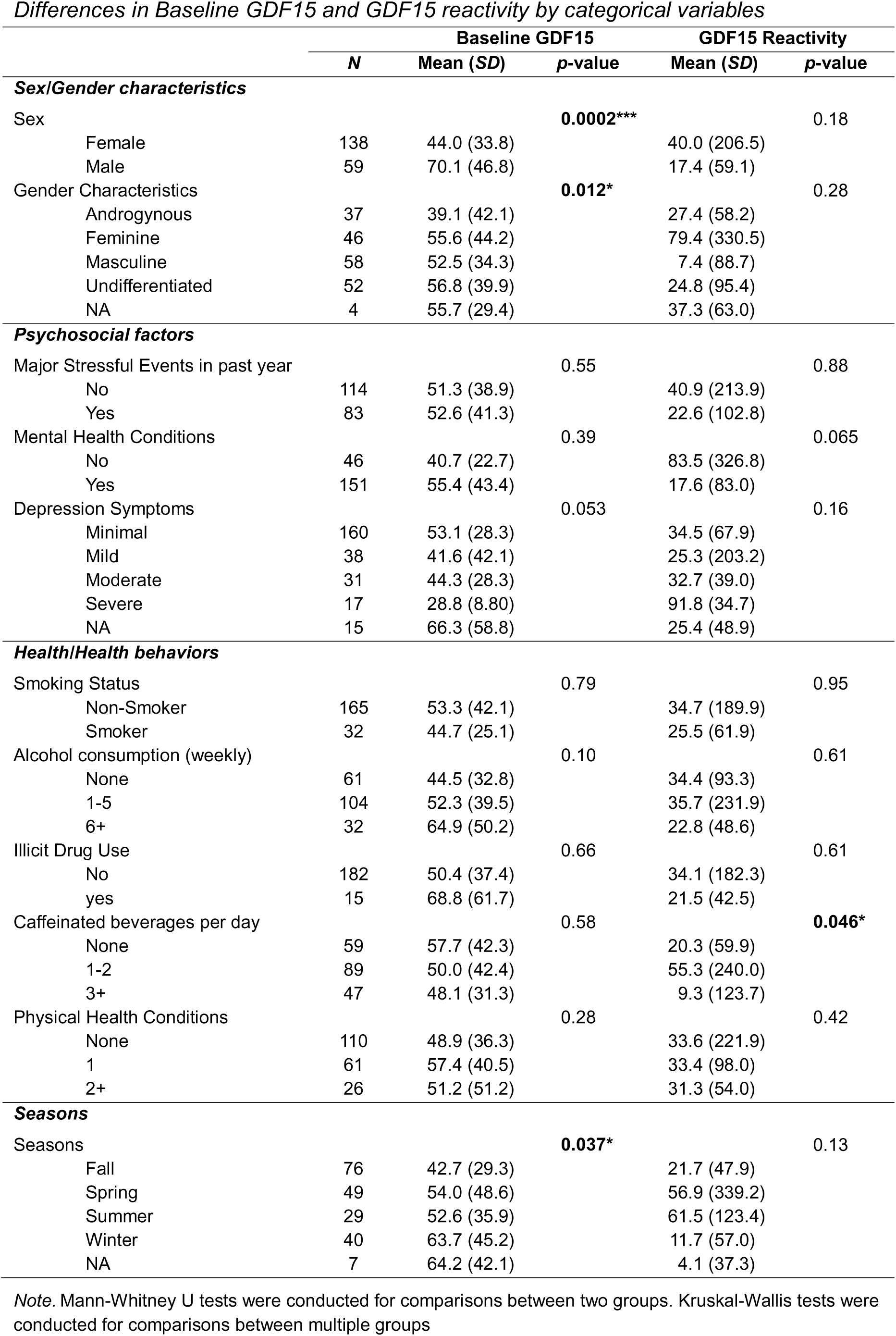
Differences in Baseline GDF15 and GDF15 reactivity by categorical variables.

|  | Baseline GDF15 |  |  | GDF15 Reactivity |  |
| --- | --- | --- | --- | --- | --- |
|  | N | Mean (SD) | p-value | Mean (SD) | p-value |
| Sex/Gender characteristics |  |  |  |  |  |
| Sex |  |  | 0.0002*** |  | 0.18 |
| Female | 138 | 44.0 (33.8) |  | 40.0 (206.5) |  |
| Male | 59 | 70.1 (46.8) |  | 17.4 (59.1) |  |
| Gender Characteristics |  |  | 0.012* |  | 0.28 |
| Androgynous | 37 | 39.1 (42.1) |  | 27.4 (58.2) |  |
| Feminine | 46 | 55.6 (44.2) |  | 79.4 (330.5) |  |
| Masculine | 58 | 52.5 (34.3) |  | 7.4 (88.7) |  |
| Undifferentiated | 52 | 56.8 (39.9) |  | 24.8 (95.4) |  |
| NA | 4 | 55.7 (29.4) |  | 37.3 (63.0) |  |
| Psychosocial factors |  |  |  |  |  |
| Major Stressful Events in past year |  |  | 0.55 |  | 0.88 |
| No | 114 | 51.3 (38.9) |  | 40.9 (213.9) |  |
| Yes | 83 | 52.6 (41.3) |  | 22.6 (102.8) |  |
| Mental Health Conditions |  |  | 0.39 |  | 0.065 |
| No | 46 | 40.7 (22.7) |  | 83.5 (326.8) |  |
| Yes | 151 | 55.4 (43.4) |  | 17.6 (83.0) |  |
| Depression Symptoms |  |  | 0.053 |  | 0.16 |
| Minimal | 160 | 53.1 (28.3) |  | 34.5 (67.9) |  |
| Mild | 38 | 41.6 (42.1) |  | 25.3 (203.2) |  |
| Moderate | 31 | 44.3 (28.3) |  | 32.7 (39.0) |  |
| Severe | 17 | 28.8 (8.80) |  | 91.8 (34.7) |  |
| NA | 15 | 66.3 (58.8) |  | 25.4 (48.9) |  |
| Health/Health behaviors |  |  |  |  |  |
| Smoking Status |  |  | 0.79 |  | 0.95 |
| Non-Smoker | 165 | 53.3 (42.1) |  | 34.7 (189.9) |  |
| Smoker | 32 | 44.7 (25.1) |  | 25.5 (61.9) |  |
| Alcohol consumption (weekly) |  |  | 0.10 |  | 0.61 |
| None | 61 | 44.5 (32.8) |  | 34.4 (93.3) |  |
| 1-5 | 104 | 52.3 (39.5) |  | 35.7 (231.9) |  |
| 6+ | 32 | 64.9 (50.2) |  | 22.8 (48.6) |  |
| Illicit Drug Use |  |  | 0.66 |  | 0.61 |
| No | 182 | 50.4 (37.4) |  | 34.1 (182.3) |  |
| yes | 15 | 68.8 (61.7) |  | 21.5 (42.5) |  |
| Caffeinated beverages per day |  |  | 0.58 |  | 0.046* |
| None | 59 | 57.7 (42.3) |  | 20.3 (59.9) |  |
| 1-2 | 89 | 50.0 (42.4) |  | 55.3 (240.0) |  |
| 3+ | 47 | 48.1 (31.3) |  | 9.3 (123.7) |  |
| Physical Health Conditions |  |  | 0.28 |  | 0.42 |
| None | 110 | 48.9 (36.3) |  | 33.6 (221.9) |  |
| 1 | 61 | 57.4 (40.5) |  | 33.4 (98.0) |  |
| 2+ | 26 | 51.2 (51.2) |  | 31.3 (54.0) |  |
| Seasons |  |  |  |  |  |
| Seasons |  |  | 0.037* |  | 0.13 |
| Fall | 76 | 42.7 (29.3) |  | 21.7 (47.9) |  |
| Spring | 49 | 54.0 (48.6) |  | 56.9 (339.2) |  |
| Summer | 29 | 52.6 (35.9) |  | 61.5 (123.4) |  |
| Winter | 40 | 63.7 (45.2) |  | 11.7 (57.0) |  |
| NA | 7 | 64.2 (42.1) |  | 4.1 (37.3) |  |

Note. Mann-Whitney U tests were conducted for comparisons between two groups. Kruskal-Wallis tests were conducted for comparisons between multiple groups

These findings build on existing literature exploring the relationship between sex hormones and blood GDF15 in the context of diseased and disordered states (Liu, Dai et al. 2019, Chen, Dai et al. 2020, Peng, Li et al. 2021, Sun, Peng et al. 2021, Li, Mei et al. 2023, Pan, Naviaux et al. 2023). Interestingly, our results align with previous studies demonstrating an inverse relationship between GDF15 and estrogen (Faubion, White et al. 2020) but contradict observations of an inverse relationship between GDF15 and testosterone (Liu, Dai et al. 2019, Chen, Dai et al. 2020, Peng, Li et al. 2021, Sun, Peng et al. 2021).

### 3.2 Salivary GDF15 and psychosocial factors

Next, we assessed the relationship between salivary GDF15 and psychosocial factors, including measures of mental health, burnout, work-related stress, chronic stress, and personality traits (**Figure 3A**). In relation to baseline salivary GDF15, we found that people who worked a greater number of hours per week had higher baseline GDF15 levels (*r_s_* = 0.17, *p* = 0.015, *p_adj_* = 0.66; **Figure 3B**).

**Figure 3.**
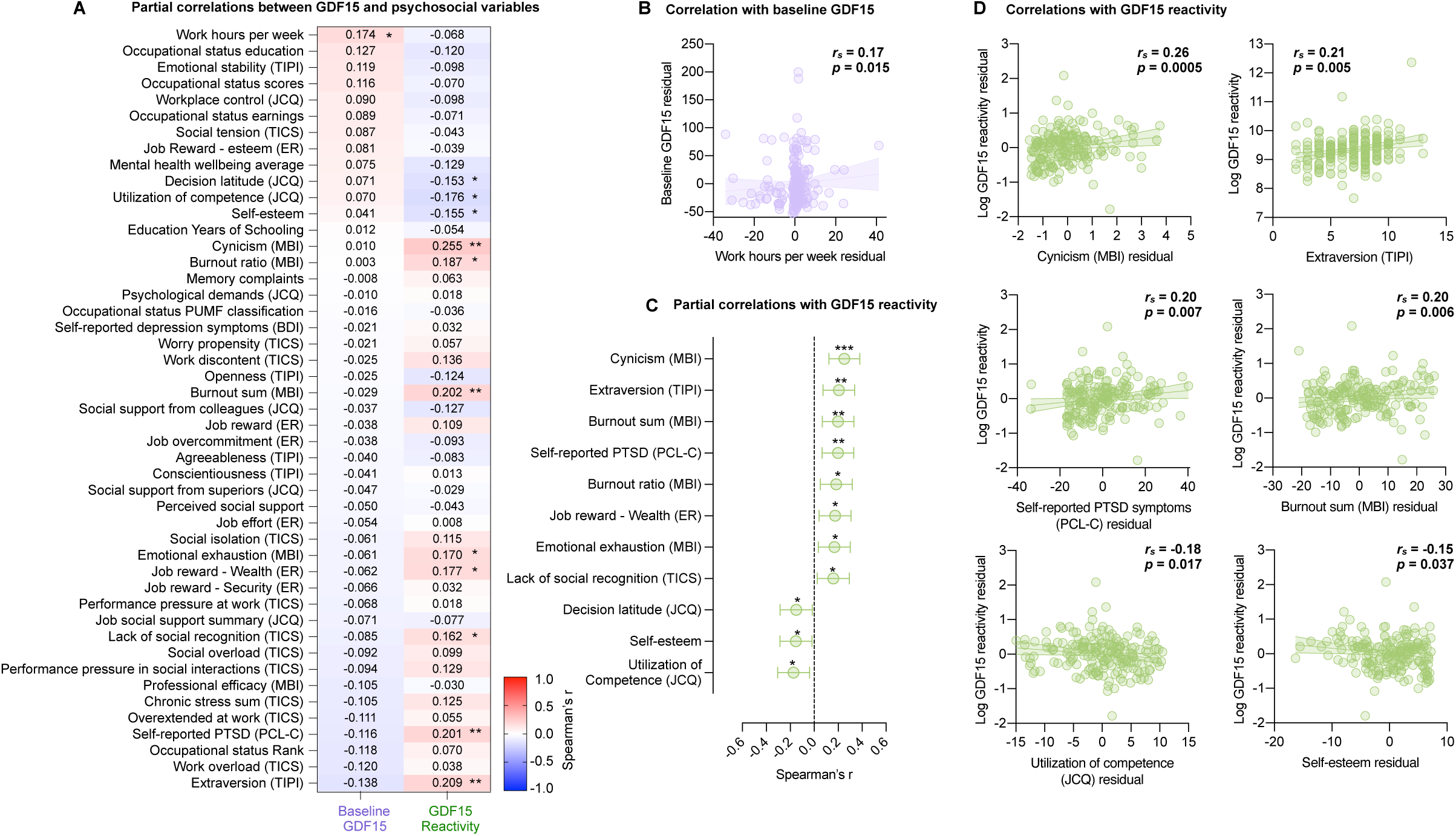
Salivary baseline GDF15 and GDF15 reactivity correlations with psychosocial factors. **(A)** Heatmap showing partial correlations between psychosocial questionnaires and baseline GDF15 or GDF15 reactivity (*n* = 156-197), adjusted for age. The heatmap was sorted by descending effect size for baseline GDF15. **(B)** Scatter plot showing the partial correlation between baseline GDF15 and work hours per week (*n* = 197) or extraversion (*n* = 186), adjusted for age. **(C)** Forest plot of significant partial correlations between GDF15 reactivity and psychosocial questionnaires (*n* = 185-186), adjusted for age. **(D)** Scatter plots showing select partial correlations between GDF15 reactivity and cynicism (*n* = 186), extraversion (*n* = 185), self-reported PTSD symptoms (*n* = 183), burnout sum (*n* = 186), utilization of competence (*n* = 186), and self-esteem (*n* = 182) scores, adjusted for age. Two-tailed significance indicated as \**p*<0.05, \*\**p*<0.01, \*\*\**p*<0.001, \*\*\*\**p*<0.0001. Unadjusted *p*-values are indicated in the figure; FDR-adjusted *p*-values can be found in Supplemental Table 1. All correlations performed using Spearman’s rank correlation.

We found greater evidence to suggest that salivary GDF15 reactivity to acute stress may be amplified in those experiencing chronic psychosocial stressors, but lower in those with more protective factors. For example, GDF15 reactivity tended to be higher in people reporting more trauma symptoms (*r_s_* = 0.20, *p* = 0.007, *p_adj_* = 0.08), work-related cynicism (*r_s_* = 0.26, *p* = 0.0005, *p_adj_* = 0.02), and emotional exhaustion (*r_s_* = 0.17, *p* = 0.021, *p_adj_* = 0.12), but lower in those who reported higher self-esteem (*r_s_* = −0.15, *p* = 0.037, *p_adj_* = 0.16), greater utilization of competence at work (*r_s_* = −0.18, *p* = 0.017, *p_adj_*= 0.11), and more work-related decision latitude (*r_s_* = −0.15, *p* = 0.038, *p_adj_*= 0.16; **Figure 3C, D**). Interestingly, GDF15 reactivity was also positively correlated with extraversion (*r_s_* = 0.21, *p* = 0.005, *p_adj_*= 0.08; **Figure 3D**), suggesting that social engagement may also influence salivary GDF15 psychobiology.

### 3.3 Salivary GDF15 and measures of health and health behaviors

Baseline GDF15 was not correlated with any measure of health or health-related behaviors, nor was it different between categories of individuals with certain behaviors or conditions (**Figure 4A, Supp. Table 1**). However, GDF15 reactivity was correlated with self-reported drug use (*r_s_*= 0.17, *p* = 0.024, *p_adj_* = 0.12) and differed according to caffeine consumption (*^2^*(3) = 6.57, *p =* 0.046). Post-hoc comparisons revealed that GDF15 was lower in those who consumed 3 or more caffeinated beverages per day compared to those who only consume 1 to 2 caffeinated beverages (*g* = 0.22, *z =* 2.36, *p* = 0.027; **Figure 4B, Supp.Table 1**); this association remained significant with outliers removed.

**Figure 4.**
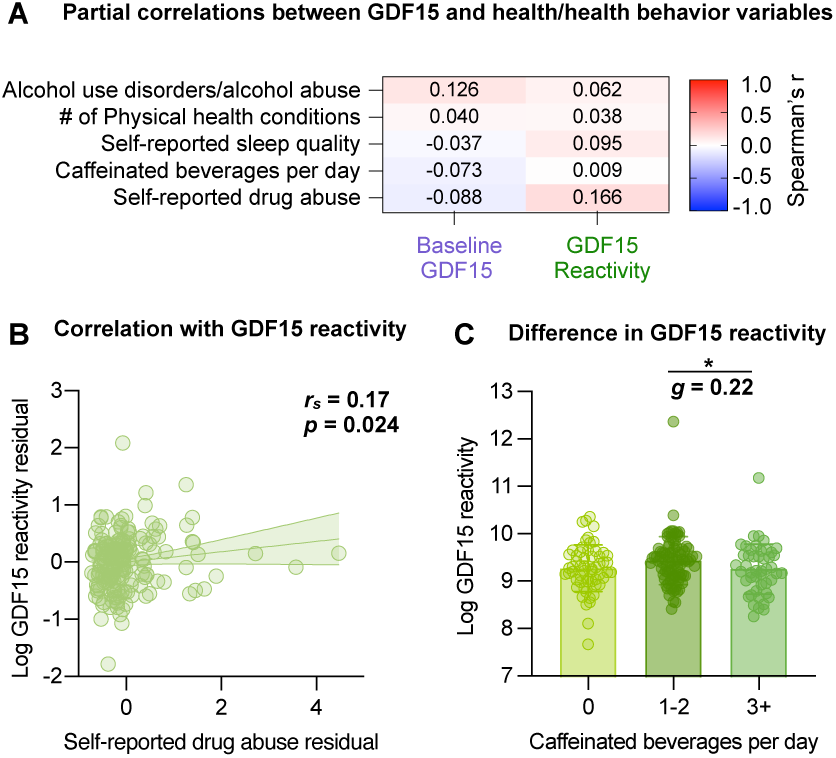
Salivary baseline GDF15 and GDF15 reactivity correlations with health and health behaviors. **(A)** Heatmap showing partial correlations between health and health behaviors and baseline GDF15 or GDF15 reactivity (*n* = 180-197), adjusted for age. The heatmap was sorted by descending effect size for baseline GDF15. Unadjusted p-values are indicated in the figure; FDR-adjusted p-values can be found in Supplemental Table 1. **(B)** Scatterplot of significant partial correlation between GDF15 reactivity self-reported drug abuse (*n* = 185), adjusted for age. **(C)** Age-adjusted log GDF15 reactivity by number of caffeinated beverages per day (0 beverages *n* = 59, 1-2 beverages *n* = 89, 3+ beverages *n* = 47). Significance of Dunn’s multiple comparison tests and Hedge’s *g* effect sizes labeled. Two-tailed significance indicated as \**p*<0.05, \*\**p*<0.01, \*\*\**p*<0.001, \*\*\*\**p*<0.0001. All correlations performed using Spearman’s rank correlation.

### 3.4 Psychosocial clustering predicts differences in GDF15

To further assess the overall pattern of relationships between salivary GDF15 and wellbeing, we used k-means clustering to categorize participants into 3 clusters based on the psychosocial and health-related behavioral variables (**Supp. Figure 1A**). Cluster 1 (*n* = 43) showed decreased age-adjusted GDF15 reactivity relative to cluster 2 (*n* = 29). There were no significant differences between cluster 3 (*n* = 46) and either of the other clusters (**Supp. Figure 1B, C**). When projected onto a principal component (PC) space, cluster 1 and 2 were most distinguished along PC1, where cluster 1 was higher than cluster 2. The top negative loadings for PC1 were variables involving psychosocial stress (e.g., burnout and chronic stress), and the top positive loadings were the composite mental health score (**Supp. Figure 1D**). This pattern establishing an axis or dimension of mental health suggests that cluster 1 participants had better overall mental health than cluster 2 participants. These results support the correlational results above that link greater psychosocial stress and fewer protective psychosocial factors to greater GDF15 reactivity.

### 3.5 Salivary GDF15 and biomarkers

We then assessed correlations between salivary GDF15 and metabolic health markers as well as allostatic load indices reflecting multisystem physiological dysregulation (Seeman, McEwen et al. 2001, Guidi, Lucente et al. 2021) (**Figure 5A**). In line with previous work connecting GDF15 with kidney disease and metabolic health (Gadd, Hillary et al. 2024, Lyu, Xv et al. 2024, Tang, Liu et al. 2024), baseline salivary GDF15 was positively correlated with creatinine (*r_s_ =* 0.29, *p* < 0.0001*, p_adj_* = 0.0005), albumin (*r_s_ =* 0.29, *p* < 0.0001*, p_adj_* = 0.0005), and waist-to-hip ratio (*r_s_ =* 0.16, *p* = 0.026*, p_adj_* = 0.16; **Figure 5B**). Salivary GDF15 reactivity was negatively correlated with albumin (*r_s_* = −0.16, *p* = 0.029, *p_adj_* = 0.56; **Figure 5C**). No significant correlations were found between GDF15 measures and allostatic load indices.

**Figure 5.**
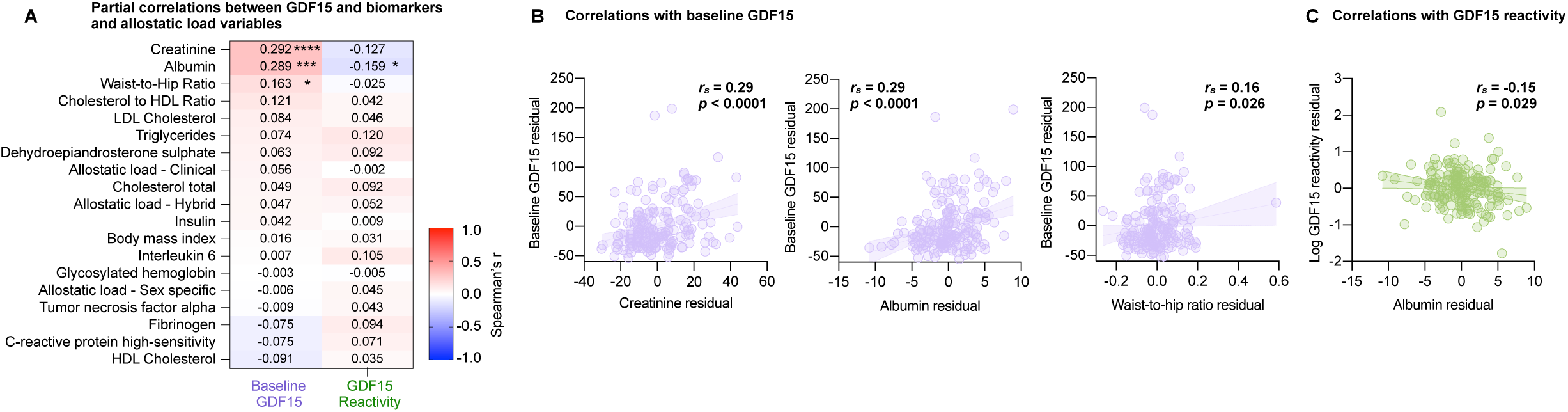
Salivary baseline GDF15 and GDF15 reactivity correlations with biomarkers and allostatic load. **(A)** Heatmap showing partial correlations between metabolic and allostatic load markers and baseline GDF15 or GDF15 reactivity (*n* = 187-197), adjusted for age. The heatmap was sorted by descending effect size for baseline GDF15. **(B)** Scatterplots of significant partial correlations between baseline GDF15 and creatinine (*n* = 192), albumin (*n* = 190), and waist-to-hip ratio (*n* = 188), adjusted for age. **(C)** Scatterplot of significant partial correlation between GDF15 reactivity albumin (*n* = 187), adjusted for age. Two-tailed significance indicated as \**p*<0.05, \*\**p*<0.01, \*\*\**p*<0.001, \*\*\*\**p*<0.0001. Unadjusted *p*-values are indicated in the figure; FDR-adjusted *p*-values can be found in Supplemental Table 1. All correlations performed using Spearman’s rank correlation.

### 3.6 Salivary GDF15 and TSST response measures

Next, we assessed whether baseline salivary GDF15 and GDF15 reactivity correlated with other parameters measured during the TSST time course, including cortisol, blood pressure, heart rate, and arterial pressure measured at different timepoints, as well as their reactivity profiles (**Figure 6A**). In comparing the time courses of GDF15 and cortisol, we found that GDF15 peaked and recovered more quickly than cortisol levels in this same cohort (**Figure 6B**). Baseline GDF15 was positively correlated with cortisol measured 40 minutes post-TSST (*r_s_* = 0.17, *p* = 0.020, *p_adj_* = 0.66; **Figure 6D**). However, GDF15 reactivity was negatively correlated with cortisol levels across several timepoints (*r_s_’s* = −0.14 to −0.24, *p’*s = 0.0007 to 0.049, *p_adj_* = 0.014 to 0.24; **Figure 6C, E**), and with the absolute (*r_s_ =* −0.23, *p =* 0.001, *p_adj_ =* 0.01) and percent (*r_s_ =* −0.19, *p =* 0.008, *p_adj_ =* 0.05) changes in cortisol from baseline to peak. Overall, this pattern of results suggest that GDF15 is a stress-responsive biomarker that may help to co-regulate cortisol reactivity, which is consistent with mouse models suggesting that GDF15 can act upstream of cortisol, signaling to the brain to trigger the hypothalamic-pituitary-adrenal axis that controls glucocorticoid secretion (Cimino, Kim et al. 2021, Ruud, Font-Gironès et al. 2024).

**Figure 6.**
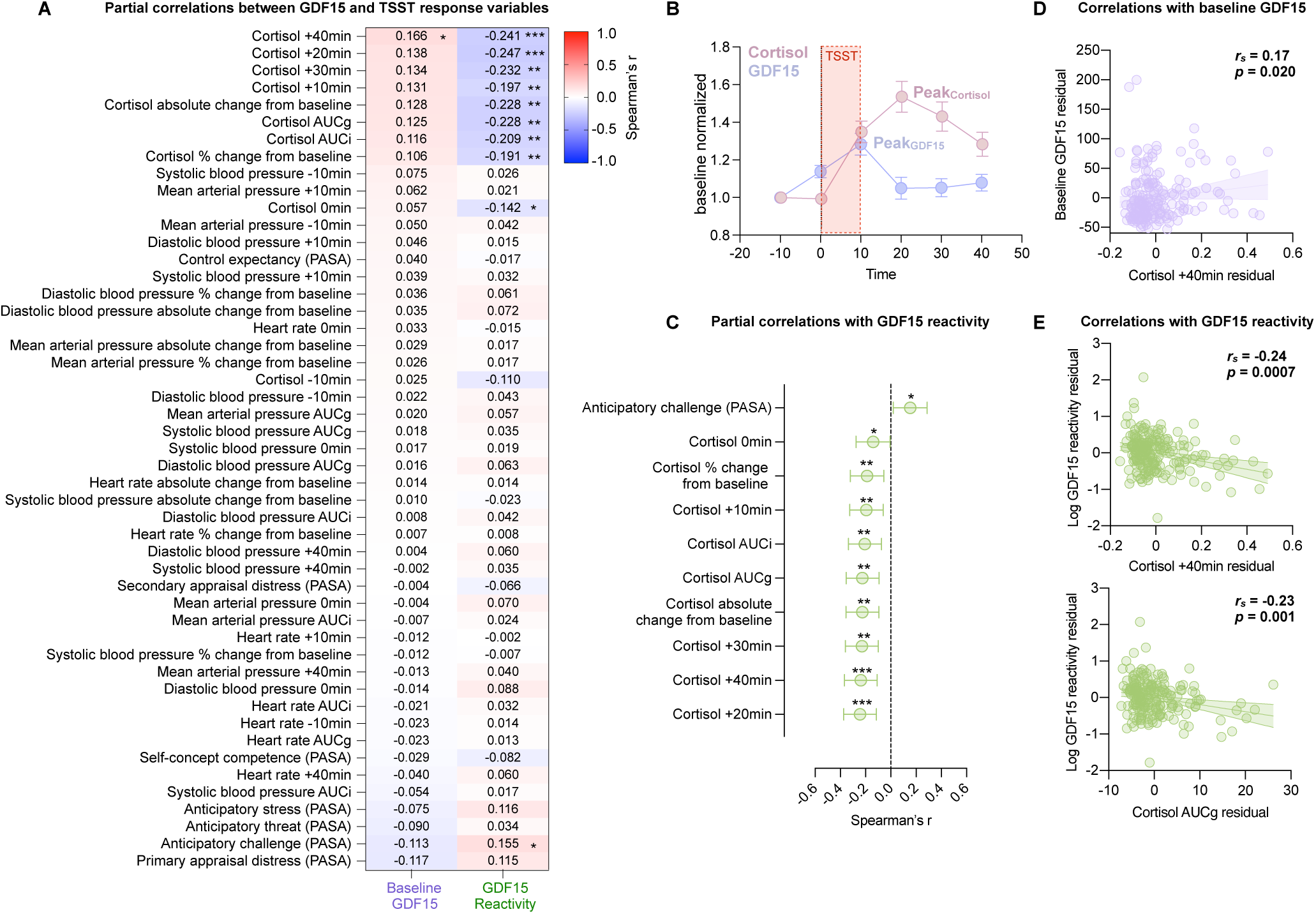
Salivary baseline GDF15 and GDF15 reactivity correlations with TSST response measures. **(A)** Heatmap showing partial correlations between TSST response measures and baseline GDF15 or GDF15 reactivity (*n* = 194-196), adjusted for age. The heatmap was sorted by descending effect size for baseline GDF15. **(B)** Average Cortisol and GDF15 time courses, normalized to baseline. Error bars represent SEM. Maximum reactivity labelled. (**C**) Forest plot of significant partial correlations between cortisol measures and GDF15 reactivity (*n* = 195), adjusted for age. **(D)** Partial correlation between baseline GDF15 and cortisol at 40 minutes post-TSST (*n* = 196), adjusted for age. **(E)** Partial correlations between log GDF15 reactivity and cortisol at 40 minutes post-TSST (*n* = 194) or cortisol Area Under the Curve calculated to ground (AUCg; *n* = 194), adjusted for age. Two-tailed significance indicated as \**p*<0.05, \*\**p*<0.01, \*\*\**p*<0.001, \*\*\*\**p*<0.0001. Unadjusted *p*-values are indicated in the figure; FDR-adjusted *p*-values can be found in Supplemental Table 1. All correlations performed using Spearman’s rank correlation.

We also note that individuals who anticipated the TSST as a challenge, rather than a threat, exhibited greater GDF15 reactivity (*r_s_ =* 0.16, *p =* 0.034, *p_adj_ =* 0.019; **Figure 6A, C**), suggesting a potential link between anticipatory stress appraisals and GDF15 regulation.

### 3.7 Considerations for using salivary GDF15 future studies

Finally, we found two additional patterns in salivary GDF15 that deserve consideration in future studies. Samples in this study were collected between October 2011 and December 2012 in Montreal, Canada. Montreal experiences harsh winters where the average temperature during the Dec 2011-Feb 2012 sample collection period varied between −23°C to 10°C (average = −9°C). Summers can also get hot, while the fall—adorned with colorful foliage—is generally considered the most comfortable season. We reasoned that if salivary GDF15 increases in response to mental and/or physical stress, the low temperatures of winter or humid heat of summer could be associated with elevated salivary GDF15. Accordingly, age-adjusted GDF15 levels did differ between seasons (*^2^*(3) = 8.46, *p =* 0.037), such that baseline GDF15 levels were lower in individuals whose saliva was collected in the fall as compared to the winter (*g* = 0.59, *Z = −*2.83, *p* = 0.014; **Figure 7A**). We found no differences in GDF15 reactivity. This seasonal variation in basal GDF15 may reflect circannual fluctuations in GDF15-inducing stressors, whether they be psychological or biological, and should be considered in future study designs and during participant recruitment.

**Figure 7.**
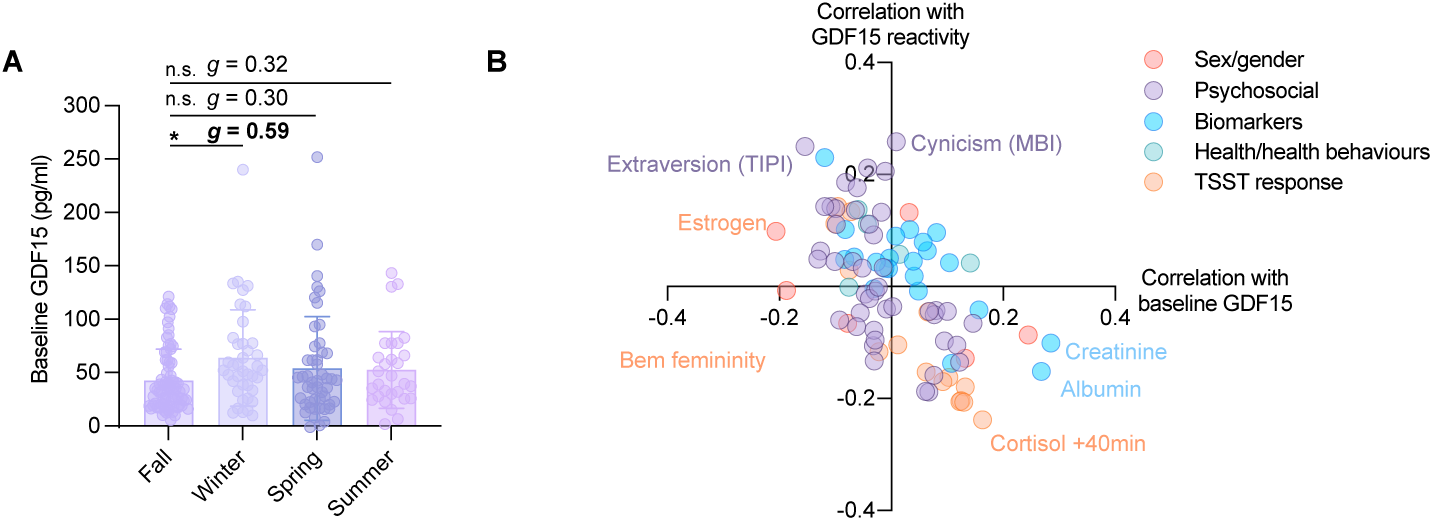
Additional patterns in salivary GDF15**(A)** Age-adjusted baseline GDF15 concentrations by season (Fall *n* = 76, Winter *n* = 40, Spring *n* = 49, Summer *n* = 29). Significance of Dunn’s multiple comparison tests and Hedge’s *g* effect sizes labeled. **(B)** Spearman’s r of partial correlations between variables and baseline GDF15 plotted against Spearman’s r partial correlations between the same variables with GDF15 reactivity. Two-tailed significance indicated as \**p*<0.05.

**Figure 8.**
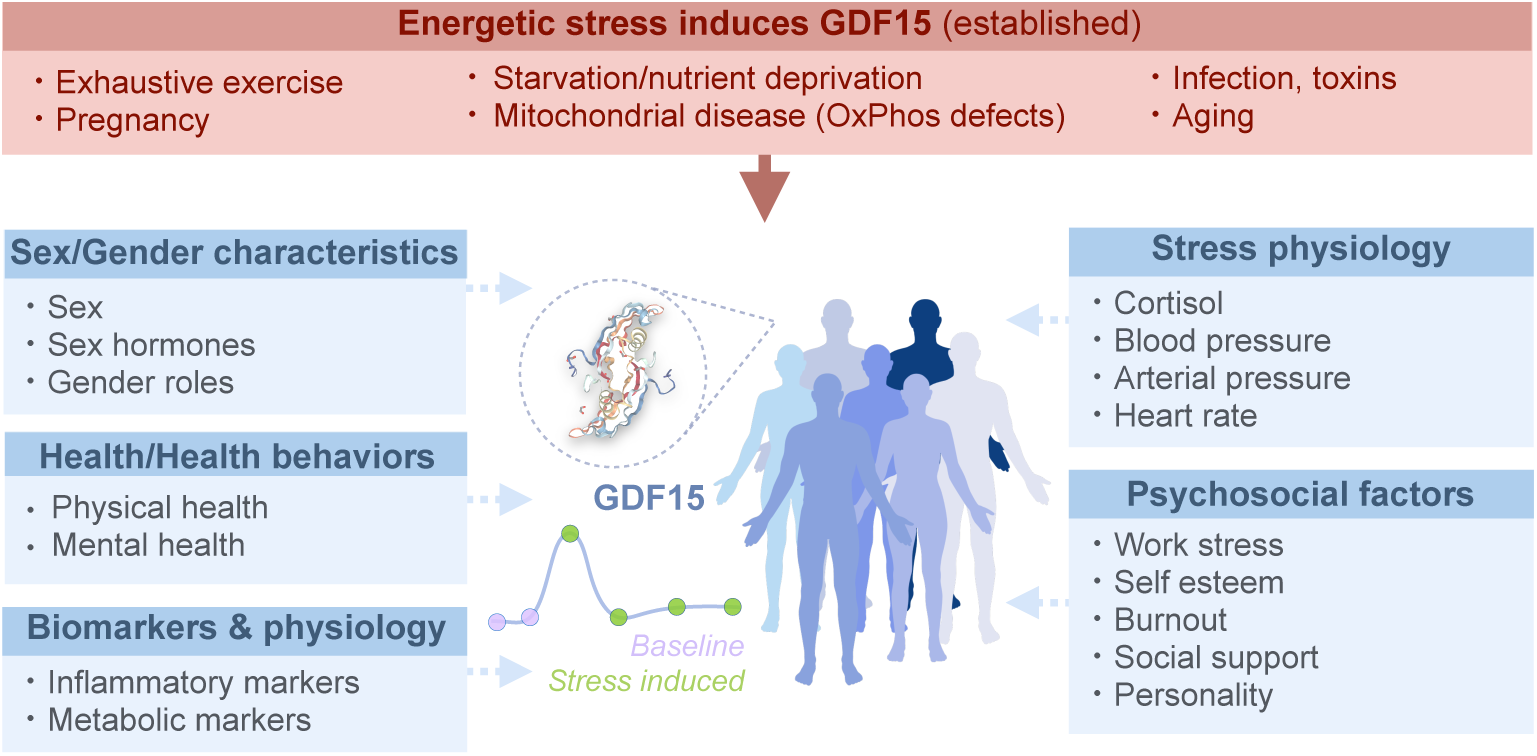
Known and unknown associations around blood and saliva GDF15. There are several known triggers and correlates of blood GDF15, many of which are energetically challenging (top). This study reports results linking baseline GDF15 and its stress reactivity with trait- and state-level biopsychosocial factors including sex and gender characteristics; measures of mental health, stress, and burnout; physical health and health behaviors; and anthropometric and blood-based metabolic biomarkers (left, and right).

The second general note relates to the comparison of baseline vs. reactivity salivary GDF15 metrics. Consistent with the negative correlation between baseline GDF15 and GDF15 reactivity (see **Figure 1D**), most correlations between these GDF15 variables and the other variables assessed were of opposite directions (**Figure 7B**).

This means that individual characteristics associated with *greater baseline* GDF15 levels tended to be associated with *lower reactivity*. Thus, researchers should consider which GDF15 metrics are best suited to their research question, as baseline and reactive GDF15 may lead to contradictory results.

## 4. Discussion

Using a well-phenotyped sample of healthy adult workers, we explored the psychobiological correlates of salivary GDF15 at baseline and reactivity over the course of a socio-evaluative stress paradigm. Consistent with recent findings from the MiSBIE study (Huang, Monzel et al. 2024), we confirm that salivary GDF15 increases acutely by an average of ∼28% with acute mental stress, positioning salivary GDF15 as a novel stress biomarker. Its biological origin positions GDF15 as a marker of energetic stress originating from mitochondria (Lockhart, Saudek et al. 2020, Monzel, Levin et al. 2024). Thus, the preliminary association patterns reported here between salivary GDF15 and a variety of psychosocial and biobehavioral factors open new opportunities for examining the shared psychobiological pathways linking human experiences and energetic stress.

In addition to these primary findings, our results warrant a few additional remarks. First, the observation that salivary GDF15 reactivity and recovery to the TSST preceded that of cortisol is consistent with the emerging psychobiological axis linking GDF15 to energy mobilization (Cimino, Kim et al. 2021, Wang, Townsend et al. 2023, Ruud, Font-Gironès et al. 2024), and potentially implicates GDF15 signaling in the regulation of canonical stress axes (e.g., the HPA axis). Second, our analyses confirmed that the association between salivary GDF15 and age is consistent with that observed using blood-based GDF15 measures. However, when comparing salivary and blood GDF15 collected from the same individual at the same time, their levels and reactivities do not correlate, suggesting that GDF15 secretion in different biofluids may be differentially regulated (Huang, Trumpff et al. 2024). Third, the novel associations between salivary GDF15 and biopsychosocial factors, including sex, traditional gender roles, trauma symptoms, work-related stress and burnout, and metabolic markers, potentially expand the spectrum of factors known to contribute to energy regulation.

Another hormone with primary effects on energy metabolism is cortisol (Picard, McEwen et al. 2018). Hypocortisolism is a marker of pathophysiology linked to fatigue, burnout, trauma, and other stress-related conditions (Fries, Hesse et al. 2005), so our results could point to the potential interaction between GDF15, cortisol, and biobehavioral experiences.

Our data also suggest the potential relevance of GDF15 in the context of mental health conditions and in the face of ongoing work stress. Although our baseline salivary GDF15 data do not show significant correlations with depression symptoms, as previously reported with blood GDF15 (Frye, Nassan et al. 2015, Mastrobattista, Lenze et al. 2023, Pan, Naviaux et al. 2023), here we present preliminary evidence showing that psychosocial stressors, including PTSD and work-related burnout, are associated with greater salivary GDF15 reactivity to acute psychosocial stress, whereas protective psychosocial factors, such as self-esteem, tend to predict less GDF15 reactivity. The positive relationships between GDF15 reactivity and extraversion and challenge appraisals of the psychosocial stress paradigm also deserve further research. These findings implicate not only objective stress or strain in salivary GDF15 regulation, but also psychological appraisal and social engagement—echoing preclinical findings linking mitochondrial biology and energy metabolism with social behaviors, motivation, and effort (Hollis, van der Kooij et al. 2015, Zalachoras, Astori et al. 2022).

### 4.1 Strengths and Limitations

In this exploratory study, we investigated the associations among salivary GDF15 and several biopsychosocial factors spanning various facets of the lived human experience. A key strength of this study is the extensive profiling of study participants and the consequent breadth of analyses, which enabled us to investigate many domains of life. However, a related limitation is the large number of comparisons performed, which largely precludes definitive conclusions around the associations of saliva GDF15 to >120 specific measures or variables (see **Supplemental Table 1**).

While additional research is needed to confirm these findings, the patterns unveiled by this study suggests that salivary GDF15 is potentially related to several clinically relevant measures of health and wellbeing. Moreover, our study sample was exclusively drawn from employees of a large psychiatric hospital, which is both a strength and limitation with respect to interpreting our work-related stress findings. The sample also constrains the generalizability of our findings to other professions. Finally, the exploratory and cross-sectional nature of this study also limits our ability to assess causality and the persistence of these relationships over time. It is our hope that the associations presented here motivate the deployment of salivary GDF15 studies to examine the energetic dimension of the human stress response, the modifiability of age-related and stress-related risk factors, and the psychobiology of well-being.

### 4.2 Conclusions

GDF15 is emerging as the most robust and specific biomarker of cellular energetic deficit currently available to researchers and clinicians (Sharma, Reinstadler et al. 2021, You, Guo et al. 2023, Deng, You et al. 2025). The inducibility of saliva GDF15 by acute mental stress and the association of GDF15 metrics with several psychosocial factors point to energetic stress as a common factor for several facets of the human experience (Bobba-Alves, Juster et al. 2022, Kelly, Trumpff et al. 2024). Figure 8 highlights known triggers for blood GDF15, together with factors explored in this study that require replication in larger studies. These results lay the foundation for future research using saliva and/or blood GDF15 to examine the energetic dimension of stress psychobiology.

## Declaration of interest

The authors have no competing interests to declare.

## Acknowledgements

The authors are grateful to Janick Boissonneault, Helen Findley, Ryan Hogan, Cécile Le Page, and Nathalie Wan, and for their support in the materials transfer. We thank Sonia J Lupien for allowing us to reuse these saliva samples for emergent discovery. We also thank Quinn Conklin for her critical advising and reading on the manuscript.

## Funding

The work of the authors was supported by NIH grants R01MH122706 and R01MH137190, the Wharton Fund, and Baszucki Group to C.T. and M.P. R.P.J. is supported by the Fonds de recherche Santé Québec and the Fondation l’Institut universitaire en santé mentale de Montréal. Data collection was supported by the Canadian Institutes of Health Research (CIHR) Institute of Gender and Health (Grant No. 222055) awarded to S. J. Lupien (Principal Investigator) while R.P.J. was a Doctoral candidate supported by the CIHR Institute of Aging (Grant No. SIA 95402) and the Research Team on Work and Mental Health (ERTSM).

## Author contributions

R.P.J. designed and coordinated the original SGA study under the supervision of Sonia, J. Lupien. C.C.L. and Q.H. performed GDF15 assays. C.C.L. analyzed the data. C.T. and M.P. advised on data analysis. C.C.L. and M.P. drafted the manuscript. C.T. and M.P. obtained funding and supervised the GDF15 analyses. All authors reviewed the final version of the manuscript.

## SUPPLEMENTAL FIGURE LEGENDS

**Figure S1.**
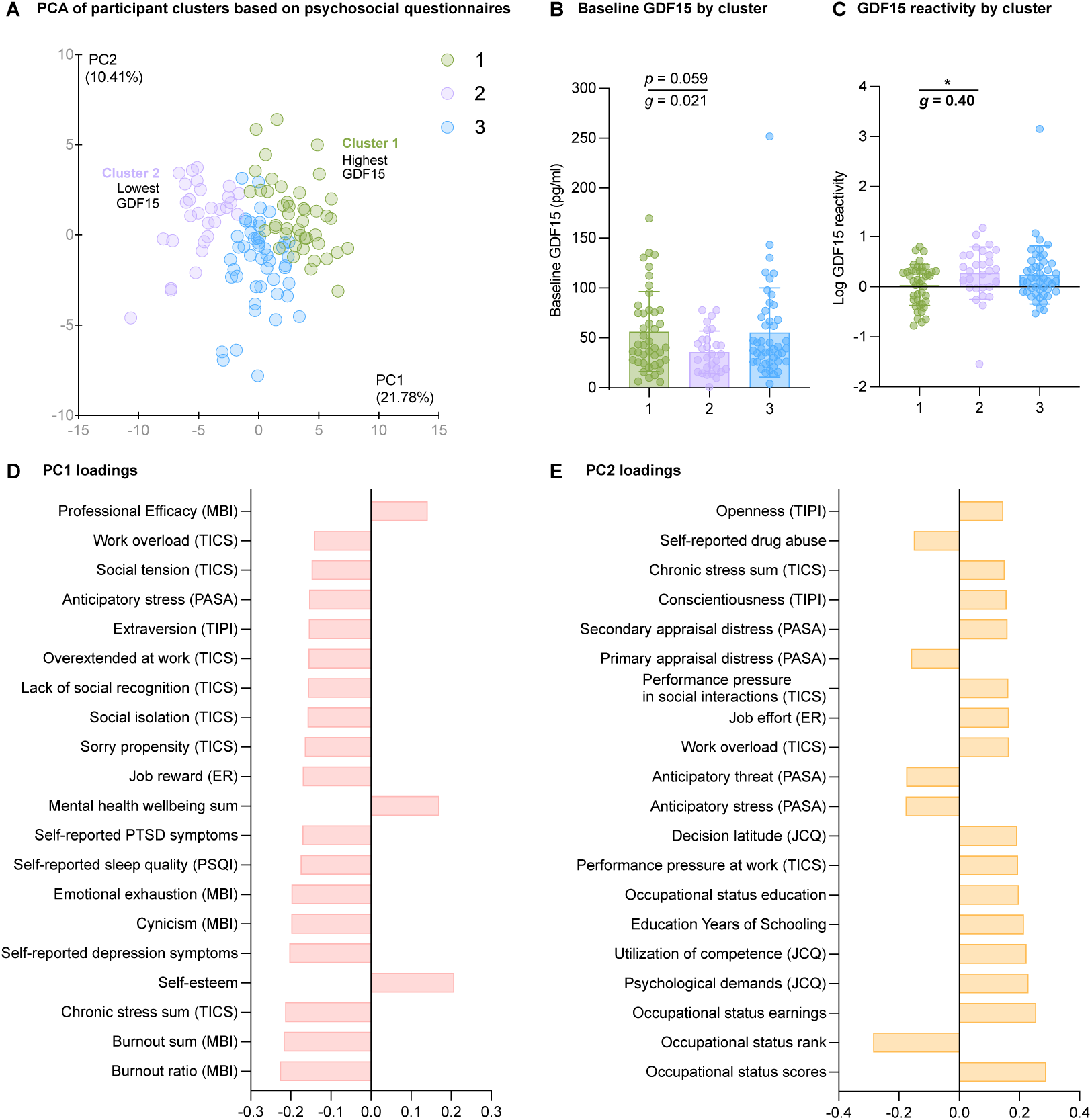
Clustering based on self-report questionnaire variables. **(A)** k-means clustering using questionnaire data, projected on a PCA space. **(B)** Age-adjusted baseline GDF15 concentrations by cluster (Cluster 1: *n* = 28, Cluster 2: *n* = 42, Cluster 3: *n* = 48). Significance of Dunn’s multiple comparison test and Hedge’s *g* effect sizes labeled. **(C)** Age-adjusted log GDF15 reactivity by clusters (Cluster 1: *n* = 28, Cluster 2: *n* = 42, Cluster 3: *n* = 48). Significance of Dunn’s multiple comparison test and Hedge’s *g* effect sizes labeled. **(D)** PC1 loadings. **(E)** PC2 loadings. Significance indicated as \**p*<0.05, \*\**p*<0.01, \*\*\**p*<0.001, \*\*\*\**p*<0.0001.

## Notes

### Competing Interest Statement

The authors have declared no competing interest.

